# Neuroanatomical Basis of Coma in Acute Ischemic Stroke

**DOI:** 10.1101/2025.08.15.670576

**Authors:** Zheng Zhang, Milo White, Jennifer A. Kim, Victor M. Torres-Lopez, Aya Khalaf, David Jin, Aaron D. Boes, Brian L. Edlow, Stephen W. English, David Fischer, Joel Geerling, Saleem M. Halablab, Maarten G. Lansberg, Margy McCullough-Hicks, James F. Meschia, Patrik Michel, Paola Palazzo, Benjamin D. Reasoner, Robert W. Regenhardt, Michael J. Young, Anh T. Tran, Seyedmehdi Payabvash, Kevin N. Sheth, Hal Blumenfeld

**Affiliations:** Department of Neurology, Yale University School of Medicine, New Haven, CT, USA; Neuroscience, Yale University School of Medicine, New Haven, CT, USA; Neurosurgery, Yale University School of Medicine, New Haven, CT, USA; Department of Neurobiology, University of Alabama at Birmingham, Birmingham, AL, USA; Departments of Neurology, Pediatrics, and Psychiatry, Carver College of Medicine, University of Iowa, Iowa City, IA, USA; Athinoula A. Martinos Center for Biomedical Imaging, Department of Radiology, Massachusetts General Hospital, Harvard Medical School, Charlestown, MA, USA; Center for Neurotechnology and Neurorecovery, Department of Neurology, Massachusetts General Hospital, Harvard Medical School, Boston, MA, USA; Department of Neurology, Mayo Clinic Florida, Jacksonville, FL, USA; Division of Neurocritical Care, Department of Neurology, Perelman School of Medicine, University of Pennsylvania, Philadelphia, PA, USA; Department of Neurology, University of Iowa, Iowa City, IA, USA; Iowa Neuroscience Institute, University of Iowa, Iowa City, IA, USA; Department of Neurology, Stanford University, Stanford, CA, USA; Department of Neurology, University of Minnesota Medical School, Minneapolis, MN, USA; The Stroke Center, Department of Neurology, Vaudois University Hospital, Lausanne University, Lausanne, Switzerland; Department of Neurosurgery, Massachusetts General Hospital, Harvard Medical School, Boston, MA, USA; Department of Radiology, Columbia University Irving Medical Center, New York, NY, USA

**Keywords:** Acute ischemic stroke, Coma, Brainstem–thalamic pathway, Lesion-symptom mapping, Subcortical arousal system, Consciousness disorders

## Abstract

**Background:** Acute ischemic stroke (AIS) can lead to profound disturbances in consciousness, including coma, which is associated with poor prognosis and increased mortality. Clarifying the lesion patterns that precipitate loss of consciousness can refine pathophysiological models and guide prognosis.

**Objectives:** In this study, we aim to identify the brain regions most commonly affected in comatose AIS and determine whether specific combinations of lesions are necessary and sufficient to produce coma.

**Methods:** We retrospectively analyzed 476 AIS patients (52 comatose) using diffusion-weighted imaging. Infarcts were automatically segmented, manually verified, and normalized to MNI space. Support vector regression lesion-symptom mapping (SVR-LSM) quantified voxel-wise associations with coma, controlling for lesion volume. To assess the necessity and sufficiency of lesion combinations, we employed permutation-based nested logistic regression models comparing all subsets of four anatomical predictors: brainstem, thalamus, cerebellum, and the rest of brain lesions.

**Results:** SVR-LSM revealed that coma was strongly associated with lesions involving the brainstem, thalamus, and cerebellum, whereas non-comatose patients exhibited predominantly cortical infarcts. Nested model comparisons showed that concurrent lesions to both the brainstem and thalamus were necessary and sufficient for coma. Additional involvement of the cerebellum or cerebral cortex did not improve predictive performance.

**Conclusions:** Coma after AIS results from a dual-node subcortical lesion pattern involving both the brainstem and thalamus. Cerebellar and cortical lesions, even when extensive, did not induce coma in the absence of the dual-brainstem and thalamic lesions. These observations emphasize the predominant role of lesion location over lesion volume in the pathogenesis of coma. They also support mechanistic models that position the brainstem and thalamic hubs as central to the neural circuitry underlying arousal. Furthermore, these findings delineate a specific anatomical substrate that may serve as a strategic target for circuit-based neuroprotective and neuromodulatory therapies.

---

Stroke is one of the leading neurological causes of severe and long-term disability worldwide. Among individuals who survive acute ischemic stroke (AIS), only approximately one-third achieve recovery with minimal or no residual deficits, whereas the majority experience moderate to severe disability that often persists for life (Slot et al., 2008). AIS is clinically defined as a sudden onset of neurological dysfunction resulting from focal brain ischemia lasting longer than 24 hours, with evidence of acute infarction on brain imaging (Sacco et al., 2013). Data from large stroke registries demonstrate that AIS can result in markedly impaired consciousness, including coma, leading to poor clinical outcomes and increased mortality (Tao et al., 2012; Vemmos et al., 2000). Lesion topography studies indicate that the infarct location, independent of lesion volume, is a primary determinant of clinical and functional outcomes. While prior work has mapped lesion patterns associated with general neurological deficits following AIS (Liu et al., 2007; Payabvash et al., 2017; Wu et al., 2015), little has been done to isolate the specific anatomical correlates of coma. Addressing this gap is essential for improving diagnostic accuracy, prognostic precision, and therapeutic strategies, and it raises important ethical considerations in the care of patients with disorders of consciousness.

Converging evidence from neuroimaging and neurophysiological studies has demonstrated that consciousness emerges from dynamic interactions between distributed cortical association networks, particularly within the prefrontal, parietal, and temporal cortices, and a deeply integrated subcortical arousal system (SAS) encompassing the basal forebrain, hypothalamus, thalamus, and upper brainstem activating systems (Blumenfeld, 2023; Fischer et al., 2016; Rohaut et al., 2019; Schiff, 2010). The SAS comprises parallel neuromodulatory pathways that regulate cortical excitability and sustain the level of arousal required for conscious processing (Saper & Scammell, 2005; Steriade & McCarley, 2005). Foundational work by Moruzzi and Magoun (1949) first identified the brainstem reticular formation as essential for wakefulness, and subsequent neuroanatomical studies have delineated an ascending arousal axis linking mesopontine nuclei to intralaminar and medial thalamic nuclei projecting to the cortex (Edlow et al., 2021, 2024; Parvizi & Damasio, 2003). Disruption of this system is associated with impaired consciousness in various clinical populations (Blumenfeld, 2012; Jang et al., 2023; Young, 2009). The mesocircuit hypothesis further provides a mechanistic model, proposing that concurrent disruption of brainstem input and central thalamic output induces frontostriatal deafferentation, globus pallidus interna disinhibition, and reduced thalamocortical drive, producing a pathological feedback loop that suppresses arousal and awareness (Schiff, 2023).

Lesion studies support the critical role of SAS structures in maintaining consciousness. Early qualitative studies reported that focal lesions in the upper brainstem, particularly the pontine tegmentum and midbrain, were strongly associated with coma (Bateman, 2001; Plum & Posner, 1982). Parvizi and Damasio (2003) further observed that in patients with brainstem strokes, lesions confined to the upper pons induced coma even in the absence of midbrain involvement. However, these early studies relied on visual assessments of lesion location rather than quantitative analyses. Advances in voxel-based lesion-symptom mapping have enabled quantitative localization of lesion-behavior relations. For example, Fischer et al. (2016) found that lesions in the rostral dorsolateral pontine tegmentum were significantly associated with coma (n = 12) compared to control patients without coma (n = 24). Yet, the small sample sizes and potential selection or reporting biases of these studies highlight the need for larger, more systematic investigations to investigate the neuroanatomical substrates of coma in AIS.

AIS presents a unique clinical model for such investigations, as it manifests brain ischemia with its complex anatomy in diverse brain regions. However, systematic mapping of coma-specific lesion sites in AIS remains unexplored. In the present study, we address this gap by using diffusion magnetic resonance imaging (MRI) data from multiple AIS repositories, applying advanced computational lesion mapping techniques not previously used for this purpose. Diffusion-weighted imaging (DWI) and the corresponding apparent diffusion coefficient (ADC) provides excellent imaging biomarkers for detecting brain ischemia in AIS. DWI and ADC demonstrate high sensitivity, ranging from 73–92% within the first three hours of stroke onset and increasing to 95–100% within six hours (Powers et al., 2019).

To sum up, the purpose of this study is to (1) identify brain areas most commonly affected in AIS with coma and to (2) determine brain lesion combinations necessary for coma in AIS. To our knowledge, this is the first study to provide direct pathological evidence linking clinical behavioral assessments of consciousness with specific lesion sites in AIS. In addition, focusing on only lesions caused by AIS rather than hemorrhagic stroke or other mass occupying brain lesions, reduces the indirect consequences of mass effect and brain distortion leading to functional impairment in brain regions outside the lesion itself. We hypothesize that coma can result from damage to bilateral subregions of the subcortical arousal systems in the brainstem and diencephalon.

## Methods

### Patients

This study utilized a multi-institutional set of retrospective datasets. Included cohorts comprise the Yale Acute Brain Injury Registry and Tissue Repository (Yale University) for both comatose and non-comatose AIS patients, and seven other sites for comatose AIS patients including the following: the Endovascular Therapy Following Imaging Evaluation for Ischemic Stroke (DEFUSE 3; Stanford University), the Iowa Neurological Patient Registry (University of Iowa Hospitals and Clinics), the Determinants of Incident Stroke Cognitive Outcomes and Vascular Effects on Recovery study (Mayo Clinic Florida), the Measuring Brain Complexity to Detect and Predict Recovery of Consciousness in the ICU study (Massachusetts General Hospital), the Functional Brain Network to Detect and Predict Consciousness Recovery study (University of Pennsylvania), and the Acute STroke Registry and Analysis of Lausanne (ASTRAL; Centre Hospitalier Universitaire Vaudois). This study was conducted with approval from the Institutional Ethics Committee of Yale University.

Eligible participants were patients with first-ever stroke, as confirmed by clinical records, who had undergone MRI scans, including DWI and ADC, within seven days of AIS onset and associated coma or for comparison, patients with AIS but no impairment of consciousness. Exclusion criteria were defined as follows: (1) Patients with intermediate levels of impaired consciousness or with causes of impaired consciousness not attributed to AIS, such as toxic-metabolic disturbances, traumatic brain injury, seizures, or the effects of intubation and/or sedative medications, were excluded. (2) Patients who were initially comatose but experienced coma reversal prior to MRI following spontaneous recovery or acute reperfusion interventions, such as intravenous thrombolysis, endovascular therapy/mechanical thrombectomy, or combined, were excluded. (3) Patients were excluded if MRI revealed major intracranial abnormalities, including previous infarcts, intracerebral hemorrhage, hemorrhagic transformation of infarction, neoplasms, abscesses, encephalitis, or traumatic injuries. Only clinically significant findings, such as confluent hemorrhages, warranted exclusion; minor petechial hemorrhages were not considered exclusionary. (4) A history of significant pre-existing neurological abnormalities, such as prior large territorial infarcts, traumatic injuries, developmental anomalies, autoimmune conditions, or neoplastic lesions, also constituted grounds for exclusion. However, age-appropriate brain atrophy and minor white matter changes consistent with chronic small vessel disease were permitted. (5) Patients with other clinically significant medical conditions known to impair consciousness were excluded, including hepatic encephalopathy, uremia, central nervous system infections (e.g., meningitis), elevated intracranial pressure, seizures within 24 hours of evaluation, and drug toxicity. (6) Patients exhibiting a midline shift greater than 5 mm on imaging were excluded, as this degree of structural displacement has been associated with impaired consciousness (Ropper, 1986; The New England Journal of Medicine). (7) Patients having a Fazekas scale score of white matter hyperintensities greater than or equal to 4 were excluded, as this has been associated with impaired cognition (Gao et al., 2021; Kynast et al., 2018; Zeng et al., 2020).

Classification of coma status or normal consciousness was based on a combination of standardized consciousness scales and detailed clinical chart review. Specifically, the Glasgow Coma Scale (GCS; Teasdale & Jennett, 1974) and the National Institutes of Health Stroke Scale (NIHSS; Kwah & Diong, 2014) item 1a were used when available. Patients who scored 4 on the GCS eye-opening item and 5 or 6 on the motor response were considered non-comatose. In contrast, patients with a GCS total score of 5 or lower—defined by scores of 1 on eye-opening and verbal response and 1 to 3 on motor response—were classified as comatose. On the NIHSS item 1a, a score of 3 was used to identify coma, while a score of 1 indicated preserved consciousness. For patients presenting in coma at stroke onset, a follow-up GCS or NIHSS item 1a was required as close to the MRI time point as possible to confirm persistent coma, particularly following any acute interventions such as thrombolysis or endovascular therapy. In cases where structured testing assessments were unavailable or considered unreliable, clinical documentation from the medical record was used to determine consciousness status based on neurological exam findings. Similarly, patients who were not in coma at presentation required a follow-up GCS or NIHSS assessment near the time of MRI, if available, to confirm their non-comatose state.

### Retrospective Datasets

#### Yale University

Imaging data were acquired using a Siemens 3.0T MAGNETOM Prisma MRI scanner. Multiband multishell DWI data were collected using a single-shot spin-echo EPI sequence (TR = 5,449 ms, TE = 95 ms, Flip angle = 90, FOV = 100 mm, slice thickness = 5 mm, matrix size = 160 × 160, 2 diffusion directions, and a maximum b-values of 1000 s/mm^2^). FLAIR images were using these parameters (TR = 9,000 ms, TE = 92 ms, flip angle = 150°, slice thickness = 5 mm, matrix size= 144 × 144, FOV = 75 mm). T1-weighted anatomical images were acquired using a MPRAGE pulse sequence (TR = 500 ms, TE = 11 ms, flip angle = 150°, slice thickness = 5 mm, matrix size = 240 × 240, FOV = 100 mm).

#### Stanford University

Imaging data were acquired using a Siemens 3.0T MAGNETOM Prisma MRI scanner. Multiband multishell DWI data were collected using a single-shot spin-echo EPI sequence (TR = 7,600 ms, TE = 72.3 ms, Flip angle = 90, FOV = 100 mm, slice thickness = 5 mm, matrix size = 128 × 128, 2 diffusion directions, and a maximum b-values of 1000 s/mm^2^). FLAIR images were using these parameters (TR = 9,500 ms, TE = 90 ms, flip angle = 160°, slice thickness = 5 mm, matrix size= 256 × 256, FOV = 100 mm). T1-weighted anatomical images were acquired using a MPRAGE pulse sequence (TR = 550 ms, TE = 55 ms, flip angle = 90°, slice thickness = 5 mm, matrix size = 192 × 192, FOV = 100 mm).

#### University of Iowa

Imaging data were acquired using a Siemens 3.0T MAGNETOM Prisma MRI scanner. Multiband multishell DWI data were collected using a single-shot spin-echo EPI sequence (TR = 3,200 ms, TE = 92 ms, Flip angle = 90°, FOV = 100 mm, slice thickness = 5 mm, matrix size = 128 × 128, 2 diffusion directions, and a maximum b-values of 1000 s/mm^2^). FLAIR images were using these parameters (TR = 9,000 ms, TE = 109 ms, flip angle = 180°, slice thickness = 5 mm, matrix size= 288 × 288, FOV = 100 mm). T1-weighted anatomical images were acquired using a MPRAGE pulse sequence (TR = 500 ms, TE = 6900 ms, flip angle = 120°, slice thickness = 1 mm, matrix size = 230 × 230, FOV = 100 mm).

#### Mayo Clinic Florida

Imaging data were acquired using a Signa HDxt 3.0T MRI scanner. Multiband multishell DWI data were collected using a single-shot spin-echo EPI sequence (TR = 1,005 ms, TE = 92 ms, Flip angle = 90°, FOV = 100 mm, slice thickness = 4 mm, matrix size = 128 × 128, 2 diffusion directions, and a maximum b-values of 1000 s/mm^2^). FLAIR images were using these parameters (TR = 1100 ms, TE = 151 ms, flip angle = 160°, slice thickness = 4 mm, matrix size= 288 × 288, FOV = 100 mm). T1-weighted anatomical images were acquired using a MPRAGE pulse sequence (TR = 483 ms, TE = 15 ms, flip angle = 90°, slice thickness = 5 mm, matrix size = 192 × 192, FOV = 100 mm).

#### Massachusetts General Hospital

Imaging data were acquired using a GE SIGNA Artist 3.0T MRI scanner. Multiband multishell DWI data were collected using a single-shot spin-echo EPI sequence (TR = 7,500 ms, TE = 104 ms, Flip angle = 90°, FOV = 100 mm, slice thickness = 5 mm, matrix size = 160 × 160, 2 diffusion directions, and a maximum b-values of 1000 s/mm^2^). FLAIR images were using these parameters (TR = 9,000 ms, TE = 91 ms, flip angle = 160°, slice thickness = 5 mm, matrix size= 224 × 224, FOV = 80 mm). T1-weighted anatomical images were acquired using a MPRAGE pulse sequence (TR = 414 ms, TE = 17 ms, flip angle = 90°, slice thickness = 5 mm, matrix size = 232 × 232, FOV = 90.625 mm).

#### University of Pennsylvania

Imaging data were acquired using a Siemens 3.0T MAGNETOM Prisma MRI scanner. Multiband multishell DWI data were collected using a single-shot spin-echo EPI sequence (TR = 3,783 ms, TE = 75 ms, Flip angle = 90°, FOV = 100 mm, slice thickness = 5 mm, matrix size = 122 × 122, 2 diffusion directions, and a maximum b-values of 1000 s/mm^2^). FLAIR images were using these parameters (TR = 11,000 ms, TE = 140 ms, flip angle = 90°, slice thickness = 5 mm, matrix size= 193 × 193, FOV = 81 mm). T1-weighted anatomical images were acquired using a MPRAGE pulse sequence (TR = 681 ms, TE = 12 ms, flip angle = 90°, slice thickness = 5 mm, matrix size = 82 × 82, FOV = 100 mm).

#### Centre Hospitalier Universitaire Vaudois

Imaging data were acquired using a Siemens 3.0T MAGNETOM MRI scanner. Multiband multishell DWI data were collected using a single-shot spin-echo EPI sequence (TR = 5,100 ms, TE = 80 ms, Flip angle = 90°, FOV = 101.37 mm, slice thickness = 1.6 mm, matrix size = 148 × 148, 2 diffusion directions, and a maximum b-values of 1000 s/mm^2^). FLAIR images were using these parameters (TR = 400 ms, TE = 2.7 ms, flip angle = 90°, slice thickness = 3 mm, matrix size= 246 × 246, FOV = 84.375 mm). T1-weighted anatomical images were acquired using a MPRAGE pulse sequence (TR = 9,000 ms, TE = 87 ms, flip angle = 150°, slice thickness = 3 mm, matrix size = 168 × 168, FOV = 75 mm).

### Brain Lesion Assessment

Diffusion-weighted image, apparent diffusion coefficient maps, and FLAIR images as Digital Imaging and Communication in Medicine (DICOM) for subjects were obtained and anonymized using DICOM Anonymizer Pro (NeoLogica s.r.l., Cairo Montenotte SV, Italy). Anonymized images were converted to the Neuroimaging Informatics Technology Initiative format (NIFTI) using dcm2nii (McCausland Center for Brain Imaging, Columbia, SC).

We used publicly available deep learning algorithms (DeepIsles) to detect and segment AIS lesions for both patients with and without coma (de la Rosa et al., 2024) using DWI, ADC, and FLAIR images. The DeepIsles achieved superior ischemic lesion detection and segmentation accuracy (median Dice score: 0.82, median lesion-wise F1 score: 0.86). Validation using a real-world external dataset (N=1686) confirmed the model’s generalizability (median Dice score: 0.82, median lesion-wise F1 score: 0.86; de la Rosa et al., 2024). All lesion masks were checked for correct segmentation by experienced neuroradiologists, blinded to the coma or non-coma state of the patients. Lesion masks were then co-registered to the Montreal Neurological Institute (MNI) space using the MNI-152 template with a 2 mm isovoxel resolution by using SPM12 (https://www.fil.ion.ucl.ac.uk/spm/software/spm12) implemented in the MATLAB environment (The MathWorks Inc., Natick, MA, USA).

## Data Analysis

### Support Vector Regression Lesion-Symptom Mapping

We applied support vector regression lesion-symptom mapping (SVR-LSM; https://github.com/atdemarco/svrlsmgui) in the MATLAB environment to identify voxel-wise associations between brain lesions and a binary clinical outcome (coma vs. non-coma). SVR-LSM is a multivariate machine learning-based approach that models all voxels simultaneously, offering advantages over traditional univariate voxel-based lesion–symptom mapping, which treats each voxel independently (DeMarco & Turkeltaub, 2018; Zhang et al., 2014). By considering the entire lesion pattern, SVR-LSM accounts for spatial correlations among voxels and is more robust to biases introduced by vascular lesion distribution (Mah et al., 2014; Sperber & Karnath, 2017). This framework reduces the likelihood of spurious associations arising from adjacent voxels and improves sensitivity to distributed effects (Pustina et al., 2018). SVR-LSM models the lesion–behavior relation using nonlinear functions, enhancing its ability to detect complex and spatially correlated patterns of lesion-deficit associations (Zhang et al., 2014). These characteristics make it particularly well-suited for studying neurological outcomes such as coma. Binarized lesion masks served as input, and each voxel was evaluated based on the presence or absence of damage across participants to estimate its contribution to the behavioral outcome. The resulting β-parameters were then remapped into three-dimensional MNI space. Voxel-wise statistical significance level in this parameter map was determined by permutation testing. Using this approach, data are randomly permuted 5,000 times, and the resulting pseudo-behavioral datasets are used to compute SVR-β maps, which represent voxel-wise weight coefficients indicating the relative contribution of each voxel’s lesion status to predicting the behavioral outcome. Finally, voxel-wise significance is determined by comparison of pseudo-behavior β-maps and the β-map obtained from real behavioral data.

Lesion volume is an important confounding variable because larger lesions tend to cause more severe deficits regardless of location (Price et al., 2017). To control for lesion volume as a confounding factor, both lesion maps and behavioral scores were residualized with respect to total lesion volume before SVR analysis (DeMarco & Turkeltaub, 2018). The outcome variable was coded categorically (coma = 1, non-coma = 0), and the hypothesis directionality was set to “high scores are bad,” ensuring that positively weighted voxels in the output map correspond to regions more predictive of coma. Because the sample sizes were imbalanced, we implemented a balanced permutation testing procedure. For each of 5,000 permutations, we randomly subsampled the non-coma group to match the number of coma cases, thereby ensuring equal group representation and minimizing bias due to class imbalance. This was implemented by modifying the internal SVR permutation code to enable stratified resampling of both behavioral labels and lesion data.

To ensure sufficient statistical power and minimize noise from sparsely lesioned voxels, analyses were restricted to voxels lesioned in at least five patients. This threshold was based on prior methodological recommendations and consistent with standards in SVR-LSM literature, which recommend a minimum overlap regardless of total sample size (Arnoux et al., 2018; Han et al., 2023). Age and sex were included as nuisance covariates. Statistical significance was assessed using 10,000 permutations. Voxel-wise results were thresholded at *p* < 0.005, and cluster-level correction for multiple comparisons was applied using a family-wise error rate of *p* < 0.05 (DeMarco & Turkeltaub, 2018). Peak coordinates of significant clusters are reported in Montreal Neurological Institute (MNI) space. Z-score maps of statistically significant voxels were generated and overlaid on anatomical templates for visualization.

### Nested Logistic Regression Modelling

To determine whether lesions restricted to individual or combined brain regions were sufficient to account for coma following AIS, we implemented a nested logistic regression modelling using the glmfit function in MATLAB R2020a, specifying a binomial distribution and logit link function. Binary lesion masks for all AIS patients (both comatose and non-comatose) were spatially normalized to the MNI-152 (2 mm) template using affine registration. Regions of interest (ROIs) were defined based on neuroanatomical structures commonly implicated in coma as identified in Aim 1. A composite anatomical atlas was constructed by integrating the Harvard Ascending Arousal Network atlas, the Morel stereotactic atlas of the thalamus, and the FreeSurfer ASEG segmentation. This multi-atlas framework provided comprehensive coverage of subcortical and infratentorial regions critically involved in consciousness regulation.

For each patient, we calculated the proportion of lesioned voxels within each ROI by dividing the number of lesioned voxels by the total number of voxels in that ROI. Additionally, we computed a residual lesion index (R), representing the proportion of lesioned voxels in the rest of the brain, i.e., the whole-brain mask excluding all defined ROIs. These lesion proportion values served as predictors in our modeling analyses. To place all predictors on a common scale and to minimize collinearity, all lesion proportion values were standardized using a *z*-transformation, in which values were centered relative to the group mean and scaled by the group standard deviation prior to model fitting. Next, we compared a series of nested logistic regression models. This approach systematically compares a sequence of nested models, where each reduced model is a proper subset of a more complex reference model. The full model included all standardized predictors, and its deviance (*D*_full_) served as the reference for evaluating nested model comparisons:

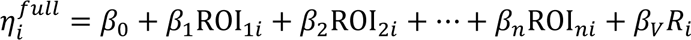

For each patient *i*, ROI_1*i*_, ROI_2*i*_, …, ROI_*ni*_ denote the regional lesion loads, and *R_i_* represents the residual lesion index. Reduced models were generated by omitting one or more predictors while retaining the others. The change in model fit was quantified as the difference in deviance (Δ*D* = *D*_reduced_ − *D*_full_), with larger Δ*D* indicating a greater loss of predictive power from excluding the corresponding ROI(s).

Statistical significance of individual regression coefficients and deviance differences was assessed through non-parametric permutation testing. Specifically, we generated 5,000 permutations in which coma labels were randomly shuffled, and both full and reduced models were refit at each iteration. The permutation p-value was computed as (k+1)/(5000+1), where k is the number of permuted Δ*D* values greater than or equal to the observed Δ*D*. Bonferroni correction was applied to control the family-wise error rate, with thresholds of α = 0.0125 for four main effects and α = 0.0167 for each model comparison. A reduced model with a non-significant permutation p-value was considered statistically equivalent to the full model, suggesting that the included regions were sufficient for predicting coma. Conversely, if all reduced models showed significantly poorer fit, this was interpreted as evidence that lesions to all regions in the full model were necessary for the occurrence of coma after AIS.

## Results

### Patients

Figure 1 presents the flowchart outlining the screening process for comatose patients, while Figure 2 illustrates the corresponding process for non-comatose patients. A total of 52 comatose and 424 non-comatose individuals were included in the study. Among the comatose cohort, the duration of coma ranged from 1.23 hours to 6 days.

**Figure 1.**
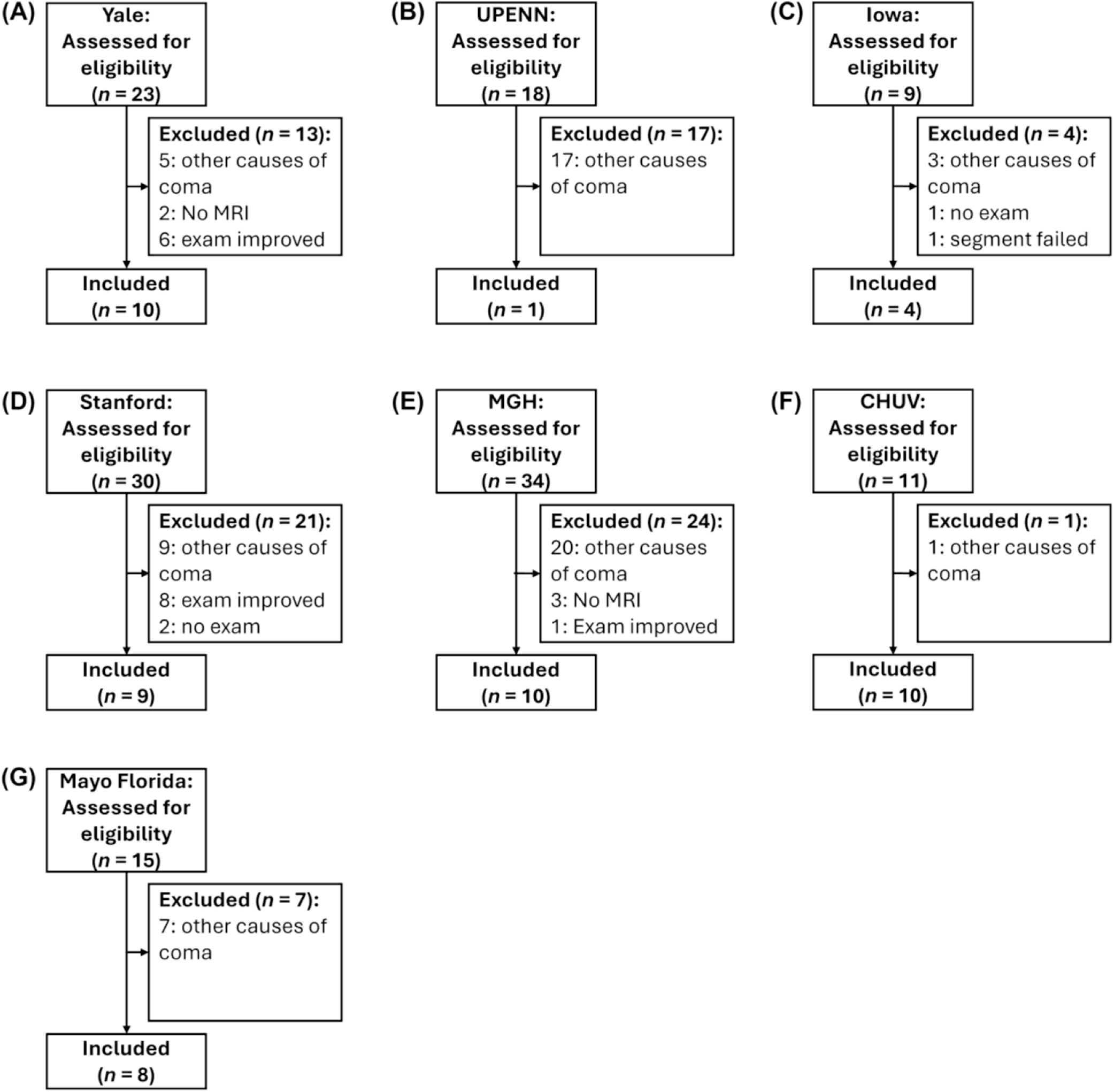
Flowchart of comatose patient screening.

**Figure 2.**
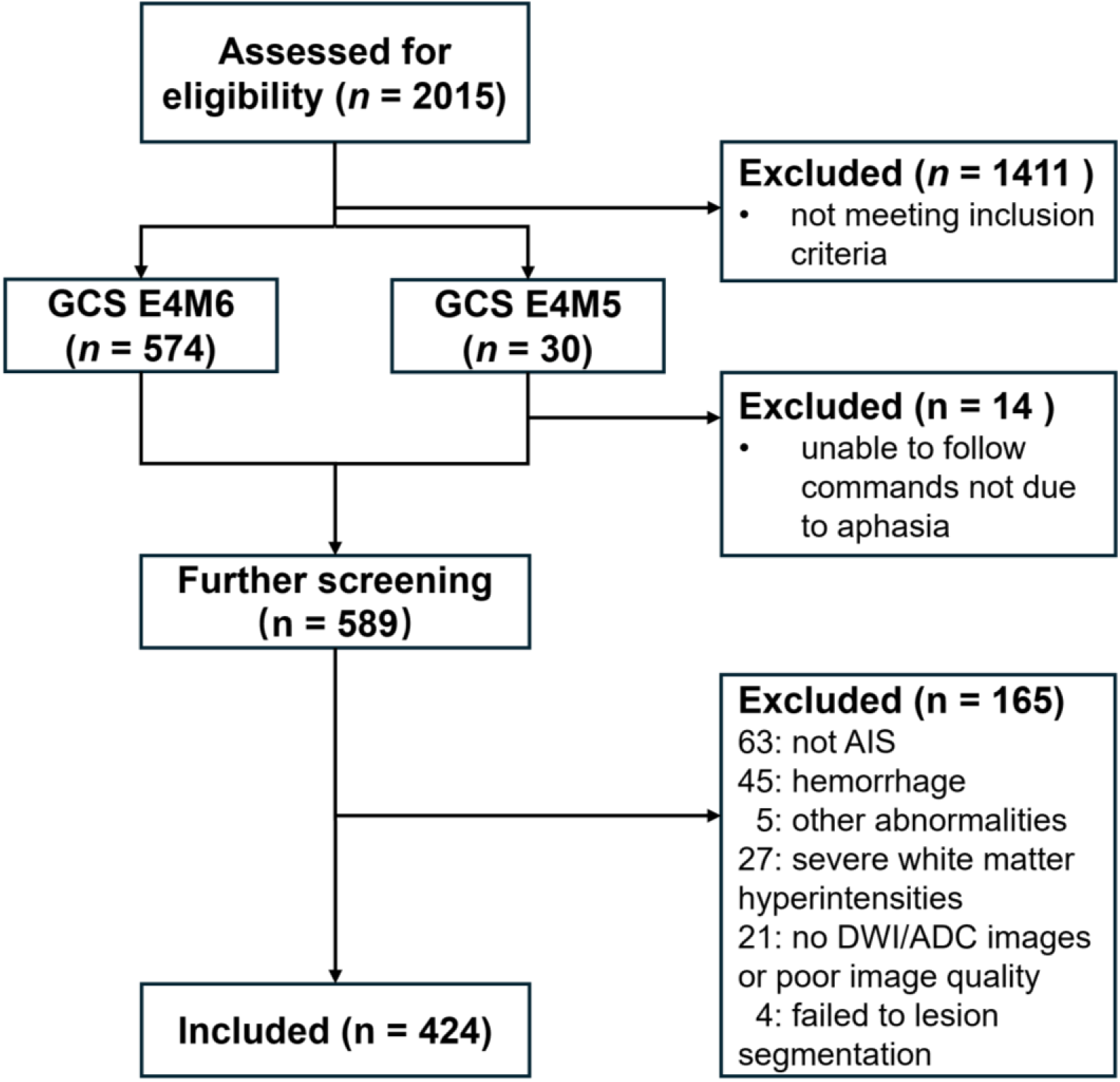
Flowchart of non-comatose patient screening. GCS, Glasgow Coma Scale; E, eye response; M, motor response; AIS, acute ischemic stroke; DWI, diffusion weighted imaging; ADC, apparent diffusion coefficient.

### Results of SVR-LSM

Figure 3A presents the overlay of lesion maps from all 476 patients, demonstrating extensive coverage across the bilateral cerebral hemispheres. The regions of maximal lesion convergence encompassed bilateral frontoparietal association cortices, temporo-occipital areas, and subcortical structures, including the brainstem, thalamus, and cerebellum. Figure 3B depicts the analysis mask, which included voxels present in at least five patients’ lesion maps (lesion overlap ≥ 5 patients).

**Figure 3.**
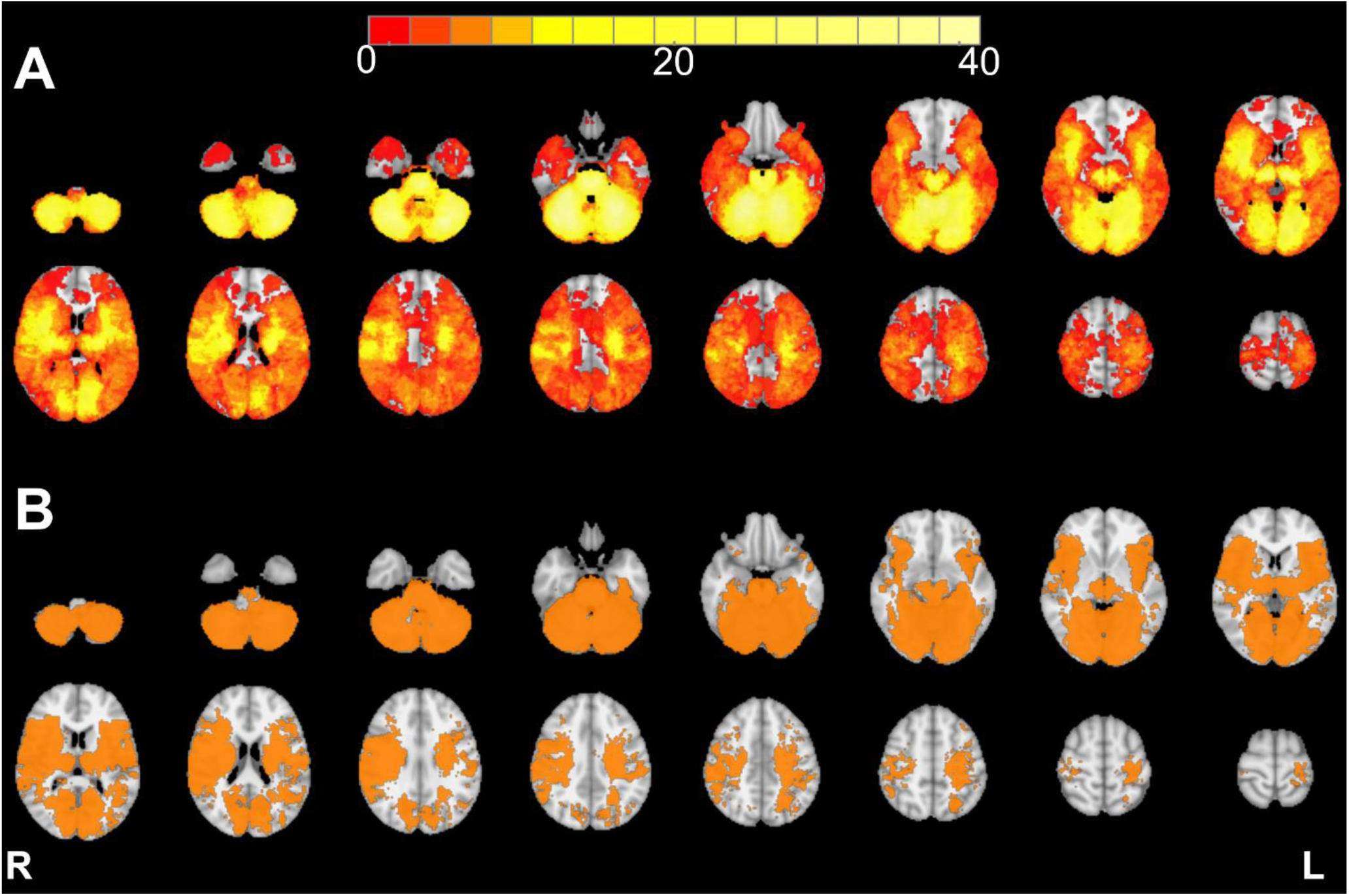
(A): Overlay images of lesions from all patients (*n* = 476). The color bar represents the number of overlapping lesion areas, with read indicating fewer overlaps and yellow indicating more overlaps. (B): Mask of voxels included in the analysis (≥ 5 in lesion overlap, shown in orange). Regions not included in this mask are excluded from the analysis.

Using SVR-LSM, we found that coma in AIS was more strongly associated with lesions involving the thalamus, brainstem, and cerebellum (Figure 4). In contrast, non-comatose patients exhibited lesion patterns predominantly localized to bilateral cortical regions, including the insula, inferior frontal gyrus, precentral gyrus, and postcentral gyrus; right hemisphere regions such as the supramarginal gyrus, Rolandic operculum, inferior parietal lobule, angular gyrus, middle frontal gyrus, and middle temporal gyrus; and left hemisphere regions including the middle occipital gyrus, superior temporal gyrus, and inferior occipital gyrus (Figure 4).

**Figure 4.**
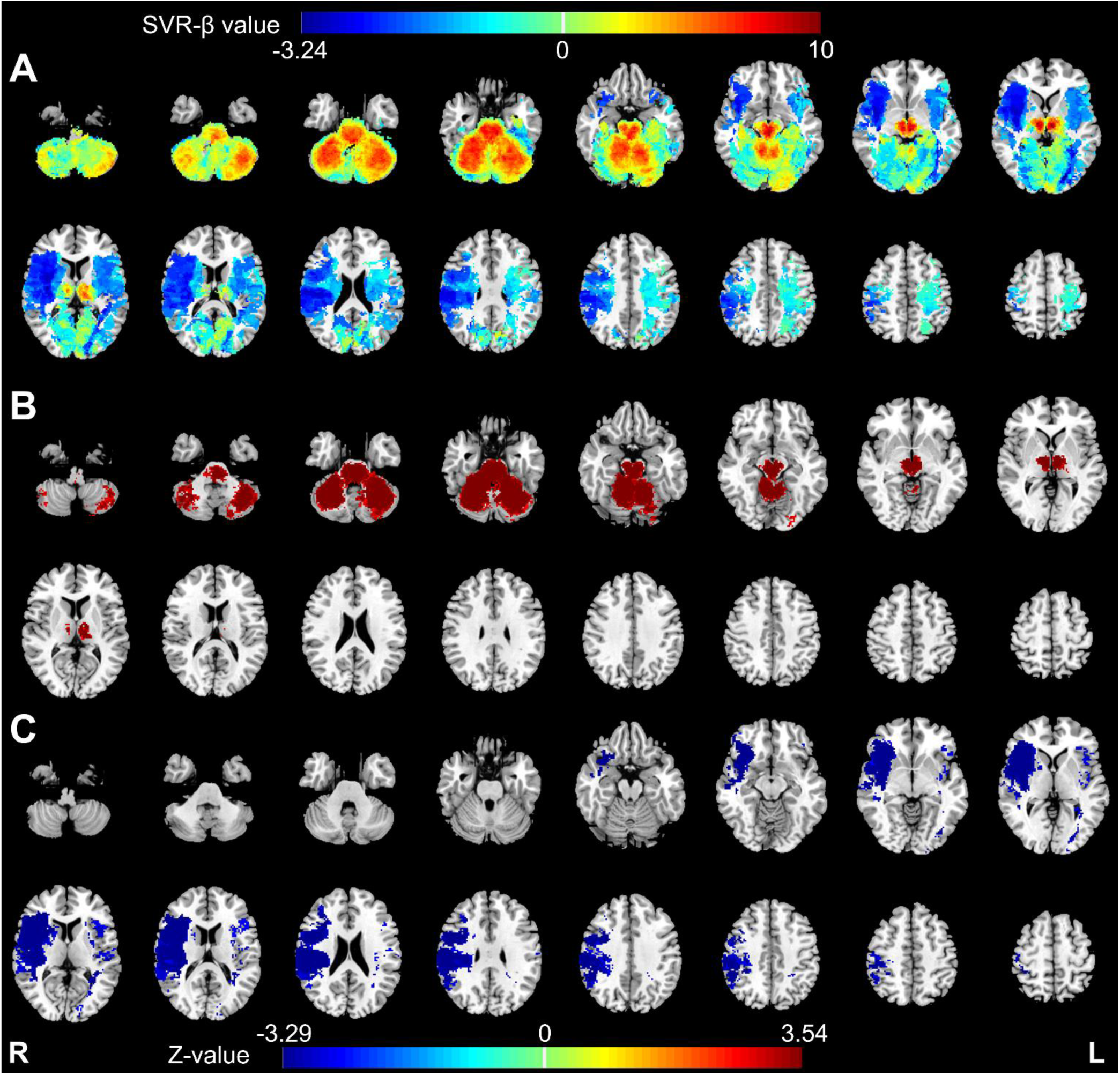
(A) Uncorrected SVR-β value output for SVR-LSM. (B & C) Parametric maps (*z*-value, voxel-wise threshold *p* < 0.005 with cluster-level corrected *p* < 0.05; 5,000 permutations) of the SVR-LSM analysis illustrate the relation between the binary lesion masks and clinical symptom (coma vs. non-coma) for all 476 patients. The regions shown in dark red indicate regions strongly associated with coma and the regions shown in blue indicate regions strongly associated with non-coma, when age and sex were used as covariates.

### Results of Nested Logistic Regression Modelling

To determine whether lesions confined to individual or paired structures could explain coma following AIS, we conducted a series of nested logistic regression analyses. Lesion proportion values within the brainstem, thalamus, and cerebellum, along with the residual lesion index, were included as predictors in the models. As shown in Table 1, logistic regression analysis of the full four-predictor model showed strong performance in explaining coma after AIS, with excellent discrimination (AUC = 0.919), acceptable calibration (Brier score = 0.095), and solid model fit (AIC = 82.47; BIC = 95.69; McFadden R² = 0.497). After Bonferroni correction, brainstem and thalamic lesions were the only significant independent predictors (β = 1.9085 and 2.5013; *p* < 0.001), while cerebellum and the residual lesion index predictors did not survive multiple comparison correction (Table 2). Excluding cerebellum and the residual lesion index predictors yielded a two-predictor model (brainstem + thalamus) with comparable fit (Δ*D* = 6.69, χ² = 0.0353) and there was no significant difference between the reduced model and the full model after Bonferroni correction (α = 0.0167; Table 3). In the brainstem + thalamus model, both retained coefficients of brainstem and thalamus remained robust (corrected *p* < 0.0005). Compared to either brainstem only model or thalamus only model, the combined brainstem + thalamus model offered significantly better fit (Δ*D* = 19.62 and 36.84, respectively, χ² < 0.0001). These findings indicate that coma risk in AIS is best explained by concurrent lesions to the brainstem and thalamus, with cerebellar or cortical involvement contributing no significant additional explanatory value.

**Table 1.**
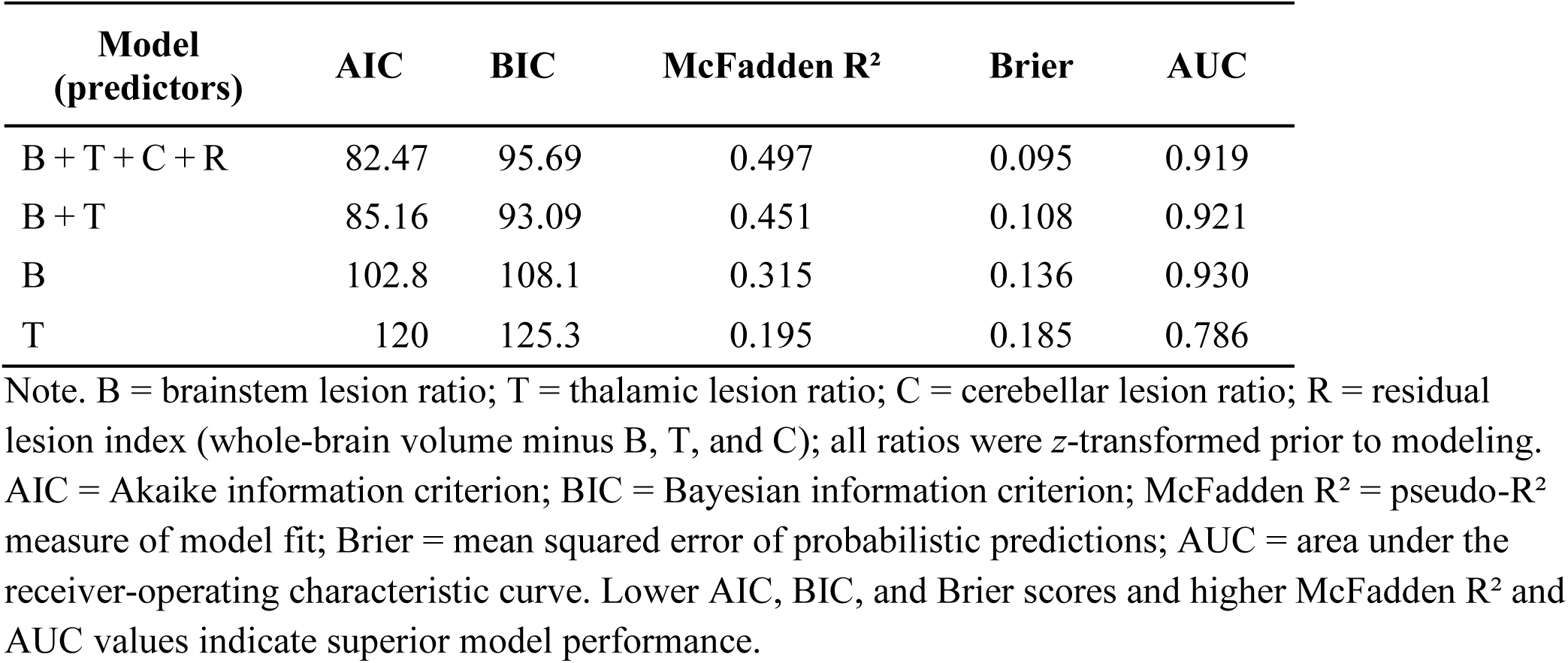
Comparative performance of nested logistic-regression models.

| **Model**<br>**(predictors)** | **AIC** | **BIC** | **McFadden R<sup>2</sup>** | **Brier** | **AUC** |
| --- | --- | --- | --- | --- | --- |
| B + T + C + R | 82.47 | 95.69 | 0.497 | 0.095 | 0.919 |
| B + T | 85.16 | 93.09 | 0.451 | 0.108 | 0.921 |
| B | 102.8 | 108.1 | 0.315 | 0.136 | 0.930 |
| T | 120 | 125.3 | 0.195 | 0.185 | 0.786 |

Note. B = brainstem lesion ratio; T = thalamic lesion ratio; C = cerebellar lesion ratio; R = residual lesion index (whole-brain volume minus B, T, and C); all ratios were z-transformed prior to modeling. AIC = Akaike information criterion; BIC = Bayesian information criterion; McFadden R<sup>2</sup> = pseudo-R<sup>2</sup> measure of model fit; Brier = mean squared error of probabilistic predictions; AUC = area under the receiver-operating characteristic curve. Lower AIC, BIC, and Brier scores and higher McFadden R<sup>2</sup> and AUC values indicate superior model performance.

**Table 2.** Permutation-based significance tests for logistic-regression models.

| Variable | Beta | SE | Perm-p | Perm-p-Bonf |
| --- | --- | --- | --- | --- |
| **Model: B + T + C + R** |  |  |  |  |
| Bonferroni-corrected $\alpha = 0.0125$ | | | | |
| Intercept | 0.8588 | 0.4074 | < 0.0005 | < 0.0005 |
| Brainstem | 1.9085 | 0.6962 | < 0.0005 | < 0.001 |
| Thalamus | 2.5013 | 0.7583 | < 0.0005 | < 0.001 |
| Cerebellum | 1.1451 | 0.5173 | < 0.005 | 0.0128 |
| Residual lesion | -0.8941 | 0.5940 | 0.0188 | 0.0752 |
| **Model: B + T** |  |  |  |  |
| Bonferroni-corrected $\alpha = 0.025$ | | | | |
| Intercept | 0.9569 | 0.4195 | < 0.0005 | < 0.0005 |
| Brainstem | 2.6027 | 0.7067 | < 0.0005 | < 0.0005 |
| Thalamus | 2.0973 | 0.6799 | < 0.0005 | < 0.0005 |
| **Model: B only** |  |  |  |  |
| Bonferroni-corrected $\alpha = 0.05$ | | | | |
| Intercept | 0.9226 | 0.4406 | < 0.0005 | < 0.0005 |
| Brainstem | 3.2679 | 0.8504 | < 0.0005 | < 0.0005 |
| **Model: T only** |  |  |  |  |
| Bonferroni-corrected $\alpha = 0.05$ | | | | |
| Intercept | 0.4872 | 0.3249 | < 0.0005 | < 0.0005 |
| Thalamus | 2.3100 | 0.6950 | < 0.0005 | < 0.0005 |

Note. B = brainstem lesion ratio; T = thalamic lesion ratio; C = cerebellar lesion ratio; R = residual
lesion index (whole-brain volume minus B, T, and C). All lesion ratios were z-standardized prior to
analysis. Perm-p values were computed from a permutation distribution of the likelihood-ratio statistic;
Perm-p-Bonf values apply a Bonferroni correction. SE = standard error of $\beta$ .

**Table 3.** Likelihood-ratio tests comparing nested logistic-regression models.

| **Reduced model** | **Deviance** | **$\Delta$ Deviance** | **$\chi^2$** |
| --- | --- | --- | --- |
| Bonferroni-corrected $\alpha = 0.0167$ | | | |
| **Reference model: B + T + C + R** | 72.47 |  |  |
| B + T | 79.16 | 6.69 | 0.0353 |
| Bonferroni-corrected $\alpha = 0.0250$ | | | |
| **Reference model: B + T** | 79.16 |  |  |
| B | 98.78 | 19.62 | < 0.0001 |
| T | 115.99 | 36.84 | < 0.0001 |

Note. B = brainstem lesion ratio; T = thalamic lesion ratio; C = cerebellar lesion ratio; R = residual
lesion index (whole-brain mask minus B, T, and C); all ratios were z-transformed prior to analysis.
Deviance is $-2 \times \log$ -likelihood. $\Delta$ Deviance equals the difference in deviance between the reduced and reference models; it follows a $\chi^2$ distribution with degrees of freedom equal to the number of constrained parameters. P-values ( $\chi^2$ ) are two-tailed and evaluated against the Bonferroni-corrected $\alpha$ thresholds reported in the table headers.

## Discussion

In this study, we employed SVR-LSM and permutation-based nested logistic regression to identify the regional determinants of coma following AIS. SVR-LSM analyses revealed that lesions involving the brainstem, thalamus, and cerebellum were significantly associated with coma, whereas non-comatose patients were more associated with lesions across a distributed set of cortical regions. Furthermore, nested model comparisons indicated that concurrent lesions to both the brainstem and thalamus were necessary and sufficient to cause coma. In contrast, cerebellar and other cerebral lesions did not contribute additional explanatory value. These findings underscore the critical importance of lesion location, particularly within core components of the SAS, as a more decisive factor than lesion volume in determining the loss of consciousness after AIS.

### Dual-Node Lesion Marker of Coma in AIS

Coma represents one of the most severe manifestations of neurological dysfunction following AI) and provides a uniquely informative model for investigating the structural substrates of consciousness. In this study, we revealed a neuroanatomical dual-node marker: coma was significantly more likely when infarcts involved both the brainstem and thalamus. In contrast, lesions restricted to either structure alone were less likely to cause coma. Moreover, once this dual-node lesion pattern was accounted for, neither total infarct volume nor additional involvement of other brain regions (e.g., the cortical areas or cerebellum) significantly improved the predictive model for coma occurrence.

Our findings align with a growing body of evidence across stroke, traumatic brain injury, and consciousness research. Clinical neuroimaging studies have shown that thalamic infarcts leading to persistent impairments in arousal often extend into the caudal midbrain tegmentum (Hindman et al., 2018), whereas isolated thalamic lesions are more frequently associated with transient reductions in consciousness, such as somnolence (Kumral et al., 2000). Complementary evidence from longitudinal diffusion MRI studies in patients with traumatic coma has demonstrated that the recovery of consciousness is accompanied by the re-establishment of structural connectivity between the brainstem and thalamus (Edlow et al., 2021; Snider et al., 2020).

Our findings are also consistent with the mesocircuit hypothesis (Schiff, 2023). In our AIS cohort, coma after AIS arises from simultaneous disruption of the brainstem and thalamus, key hubs essential for sustaining corticothalamic activity and wakefulness (Edlow et al., 2021). Infarcts affecting the paramedian midbrain or pons interrupt ascending neuromodulatory input, leading to hyperpolarization of thalamic relay neurons. Concurrent thalamic lesion further impairs thalamocortical output and striatal excitation, resulting in excessive pallidal inhibition, widespread cortical hypometabolism, and functional network disconnection (Saper et al., 2005). Therapeutic approaches targeting these circuits, such as deep brain stimulation or focused ultrasound of the central lateral thalamus, have shown potential to restore effective arousal dynamics and, in some cases, consciousness. These findings support a refined mesocircuit model in which coma reflects a strategic subcortical disconnection involving both ascending brainstem-to-thalamus and thalamocortical pathways.

Collectively, our findings reinforce the central role of the brainstem–thalamic circuit as a critical pathway for the maintenance of consciousness, providing large-scale, lesion-based evidence in AIS that coma is most probable when both the brainstem and thalamus are concurrently compromised. This dual-node model highlights that the pathogenesis of coma is predominantly driven by lesions in functionally critical subcortical structures essential for consciousness, rather than by lesion volume, thereby advancing our understanding of the key neuroanatomical substrates underpinning the maintenance of consciousness.

### The Cerebellum and Associative Loops

Although the cerebellum has been increasingly implicated in higher-order cognitive and affective functions via dentato-thalamo-cortical pathways (Dum & Strick, 2003; Stoodley & Schmahmann, 2018), our findings suggest that cerebellar integrity is neither necessary nor sufficient for the maintenance of consciousness after AIS. In permutation-based nested logistic regression analyses, cerebellar lesions did not significantly improve the prediction of coma once brainstem and thalamic involvement were accounted for. This lack of statistical contribution is biologically plausible: dentato-thalamo-cortical projections converge on thalamic relay nuclei that were already structurally compromised in comatose patients. When these thalamic targets are damaged, cerebellar output lacks an effective route to the cortex, rendering additional cerebellar injury insufficient to further impair arousal (Pelzer et al., 2017). Clinically, isolated cerebellar infarcts cause motor deficits such as dysmetria and ataxic dysarthria but preserve full alertness (Schmahmann, 2019). Consistent with these prior studies, coma in our AIS cohort occurred only when cerebellar lesions co-occurred with brainstem–thalamic damage, underscoring the cerebellum’s modulatory, rather than primary, role in sustaining consciousness.

### Limitations and Implications

Two limitations warrant consideration. First, hyperacute diffusion-weighted imaging does not capture dynamic pathological processes such as peri-infarct edema, hemorrhagic transformation, or diaschisis, which may influence clinical outcomes. Second, our analyses did not account for white matter disconnection, a critical factor in understanding how focal lesions propagate dysfunction across large-scale brain networks and contribute to consciousness impairment.

Our findings emphasize the clinical significance of early MRI identification of combined brainstem and thalamic lesions to inform proactive management and accurate prognostication. Crucially, preserved subcortical hubs can sustain arousal despite extensive cortical damage, underscoring the limitations of prognosis based solely on cortical injury. From a therapeutic perspective, neuromodulation trials, such as central thalamic DBS, focused ultrasound, and closed-loop vagus nerve stimulation, should stratify AIS patients according to residual brainstem–thalamic integrity using structural and functional connectomic biomarkers (Edlow et al., 2021; Warren et al., 202;). Moreover, closed-loop systems guided by EEG arousal signatures may optimize stimulation parameters and enhance outcomes (Hebron et al., 2024). Integrating these biomarkers into trial design will improve patient selection and maximize the efficacy of circuit-level interventions.

Future research employing high-angular-resolution diffusion imaging alongside advanced disconnectome mapping could determine whether partial preservation of SAS pathways (e.g., medial forebrain bundle, tegmento-thalamic tracts) supports wakefulness despite focal hub damage (Bruno et al., 2023). Longitudinal imaging would enable tracking of microstructural changes from acute to chronic stages (Alegiani et al., 2019; Park et al., 2022; Wang et al., 2019). Concurrent high-density EEG and resting-state fMRI, temporally aligned, may elucidate how disconnection disrupts thalamo-cortical synchrony, workspace ignition, and network dynamics (Demertzi et al., 2019). These integrative methods could clarify the progression from focal lesions to global dysfunction and identify electrophysiological markers predictive of arousal collapse (Soddu et al., 2020)

## Conclusion

This study identifies concurrent lesions to the brainstem and thalamus as the key determinants of coma after AIS, highlighting lesion location within the subcortical arousal system as more critical than lesion volume or cerebellar involvement. Early detection of combined brainstem–thalamic damage should guide clinical management and prognostication. These findings support targeted, circuit-based therapies focused on preserving or restoring these core hubs to improve recovery of consciousness after stroke.

## Acknowledgements

This work was supported by American Heart Association (24POST1195051) https://doi.org/10.58275/AHA.24POST1195051.pc.gr.190774

## References

Alegiani, A. C., MacLean, S., Braass, H., et al. (2019). Dynamics of water diffusion changes in different tissue compartments from acute to chronic stroke—A serial diffusion tensor imaging study. Frontiers in Neurology, 10, 158. 10.3389/fneur.2019.00158

Arnoux, A., Toba, M. N., Duering, M., Diouf, M., Daouk, J., Constans, J. M., … & Godefroy, O. (2018). Is VLSM a valid tool for determining the functional anatomy of the brain? Usefulness of additional Bayesian network analysis. Neuropsychologia, 121, 69–78. 10.1016/j.neuropsychologia.2018.10.003

Bateman, D. E. (2001). Neurological assessment of coma. *Journal of Neurology*, Neurosurgery & Psychiatry, 71(suppl 1), i13–i17. 10.1136/jnnp.71.suppl_1.i13

Blumenfeld, H. (2012). Impaired consciousness in epilepsy. The Lancet Neurology, 11(9), 814–826. 10.1016/S1474-4422(12)70188-6

Blumenfeld, H. (2023). Brain mechanisms of conscious awareness: Detect, pulse, switch, and wave. The Neuroscientist, 29(1), 9–18. 10.1177/10738584221132020

Bruno, M.-A., Gosseries, O., Demertzi, A., Vanhaudenhuyse, A., Thibaut, A., & Laureys, S. (2023). Disorders of consciousness: Moving from passive diagnosis to active neurostimulation. Nature Reviews Neurology, 19(3), 183–198. 10.1038/s41582-023-00764-7

de la Rosa, E., et al. (2024). A robust ensemble algorithm for ischemic stroke lesion segmentation: Generalizability and clinical utility beyond the ISLES challenge. arXiv, 2403.19425. 10.48550/arXiv.2403.1942

DeMarco, A. T., & Turkeltaub, P. E. (2018). A multivariate lesion symptom mapping toolbox and examination of lesion-volume biases and correction methods in lesion-symptom mapping (Vol. 39, No. 11, pp. 4169–4182). Hoboken, USA: John Wiley & Sons, Inc.

Demertzi, A., Tagliazucchi, E., Dehaene, S., Deco, G., Barttfeld, P., Raimondo, F., … Laureys, S. (2019). Human consciousness is supported by dynamic complex patterns of brain signal coordination. Science Advances, 5(2), eaat7603. 10.1126/sciadv.aat7603

Dum, R. P., & Strick, P. L. (2003). An unfolded map of the cerebellar dentate nucleus and its projections to the cerebral cortex. Journal of Neurophysiology, 89(1), 634–639. 10.1152/jn.00626.2002

Edlow, B. L., Chatelle, C., Spencer, C. A., … Fischer, D. B. (2021). Therapies to restore consciousness after brain injury: A gap analysis and future directions. Neurocritical Care, 35(Suppl 1), 68–85. 10.1007/s12028-021-01227-y

Edlow, B. L., Olchanyi, M., Freeman, H. J., et al. (2024). Multimodal MRI reveals brainstem connections that sustain wakefulness in human consciousness. Science Translational Medicine, 16(745), eadj4303. 10.1126/scitranslmed.adj4303

Fischer, D. B., Boes, A. D., Demertzi, A., Evrard, H. C., Laureys, S., Edlow, B. L., … & Geerling, J. C. (2016). A human brain network derived from coma-causing brainstem lesions. Neurology, 87(23), 2427–2434. 10.1212/WNL.0000000000003404

Gao, Y., Gu, H., Yang, J., Li, Z., & Zhang, J. (2021). Prognosis of patients in prolonged coma after severe carbon monoxide poisoning. Human & Experimental Toxicology, 40(8), 1355–1361. 10.1177/0960327121997992

Han, I. J., Lee, J. E., Song, H. N., Baek, I. Y., Choi, J., Chung, J. W., … & Seo, W. K. (2023). Imaging and clinical predictors of acute constipation in patients with acute ischemic stroke. Frontiers in Neuroscience, 17, 1263693. 10.3389/fnins.2023.1263693

Hebron, H., Lugli, B., Dimitrova, R., Jaramillo, V., Yeh, L. R., Rhodes, E., … Violante, I. R. (2024). A closed-loop auditory stimulation approach selectively modulates alpha oscillations and sleep onset dynamics in humans. PLOS Biology, 22(6), e3002651. 10.1371/journal.pbio.3002651

Jang, S. H., Kwak, S., & Lee, M. Y. (2023). Prognosis prediction for impaired consciousness recovery in stroke patients using videofluoroscopic swallowing study: A retrospective observational study. Medicine, 102(20), e33860. 10.1097/MD.0000000000033860

Kumral, E., Evyapan, D., & Kutluhan, S. (2000). Pure thalamic infarctions: clinical findings. Journal of Stroke and Cerebrovascular Diseases, 9(6), 287–297. 10.1053/jscd.2000.18741

Kwah, K. B., & Diong, J. H. (2014). National Institutes of Health Stroke Scale item 1a predicts consciousness level in acute stroke. Journal of Clinical Neuroscience, 21(9), 1657–1659. 10.1016/j.jocn.2013.12.032

Kynast, J., Lampe, L., Luck, T., Frisch, S., Arelin, K., Hoffmann, K. T., … & Schroeter, M. L. (2018). White matter hyperintensities associated with small vessel disease impair social cognition beside attention and memory. Journal of Cerebral Blood Flow & Metabolism, 38(6), 996–1009. 10.1177/0271678X1771938

Liu, Y., Karonen, J. O., Nuutinen, J., Vanninen, E., Kuikka, J. T., & Vanninen, R. L. (2007). Crossed cerebellar diaschisis in acute ischemic stroke: A study with serial SPECT and MRI. Journal of Cerebral Blood Flow & Metabolism, 27(10), 1724–1732. 10.1038/sj.jcbfm.9600467

Mah, Y. H., Husain, M., Rees, G., & Nachev, P. (2014). Human brain lesion-deficit inference remapped. Brain, 137(9), 2522–2531. 10.1093/brain/awu164

Mashour, G. A., Roelfsema, P., Changeux, J. P., & Dehaene, S. (2020). Conscious processing and the global neuronal workspace hypothesis. Neuron, 105(5), 776–798. 10.1016/j.neuron.2020.01.026

Moruzzi, G., & Magoun, H. W. (1949). Brain stem reticular formation and activation of the EEG. Electroencephalography and Clinical Neurophysiology, 1(1–4), 455–473. 10.1016/0013-4694(49)90219-9

Park, M., Cho, Y., Kim, D. H., et al. (2022). Myelin water imaging of nerve recovery in rehabilitating stroke patients. Journal of Magnetic Resonance Imaging, 56(5), 1548–1556. 10.1002/jmri.28185

Parvizi, J., & Damasio, A. R. (2003). Neuroanatomical correlates of brainstem coma. Brain, 126(7), 1524–1536. 10.1093/brain/awg166

Payabvash, S., Taleb, S., Benson, J. C., & McKinney, A. M. (2017). Acute ischemic stroke infarct topology: Association with lesion volume and severity of symptoms at admission and discharge. American Journal of Neuroradiology, 38(1), 58–63. 10.3174/ajnr.A4970

Pelzer, E. A., Melzer, C., Timmermann, L., von Cramon, D. Y., & Tittgemeyer, M. (2017). Basal ganglia and cerebellar interconnectivity within the human thalamus. Brain Structure and Function, 222(1), 381–392. 10.1007/s00429-016-1223-z

Plum, F., & Posner, J. B. (1982). The diagnosis of stupor and coma (Vol. 19). Oxford University Press, USA.

Powers, W. J., Rabinstein, A. A., Ackerson, T., Adeoye, O. M., Bambakidis, N. C., Becker, K., … & Tirschwell, D. L. (2019). Guidelines for the early management of patients with acute ischemic stroke: 2019 update to the 2018 guidelines for the early management of acute ischemic stroke: a guideline for healthcare professionals from the American Heart Association/American Stroke Association. Stroke, 50, 12. 10.1161/STR.0000000000000211

Price, C. J., Hope, T. M., & Seghier, M. L. (2017). Ten problems and solutions when predicting individual outcome from lesion site after stroke. NeuroImage, 145, 200–208. 10.1016/j.neuroimage.2016.08.00

Pustina, D., Avants, B., Faseyitan, O. K., Medaglia, J. D., & Coslett, H. B. (2018). Improved accuracy of lesion to symptom mapping with multivariate sparse canonical correlations. Neuropsychologia, 115, 154–166. 10.1016/j.neuropsychologia.2017.08.027

Rohaut, B., Doyle, K. W., Reynolds, A. S., Igwe, K., Couch, C., Matory, A., … & Claassen, J. (2019). Deep structural brain lesions associated with consciousness impairment early after hemorrhagic stroke. Scientific Reports, 9(1), 4174. 10.1038/s41598-019-41042-2

Sacco, R. L., Kasner, S. E., Broderick, J. P., et al. (2013). An updated definition of stroke for the 21st century: A statement for healthcare professionals from the American Heart Association/American Stroke Association. Stroke, 44(7), 2064–2089. 10.1161/STR.0b013e318296aeca

Saper, C. B., Scammell, T. E., & Lu, J. (2005). Hypothalamic regulation of sleep and circadian rhythms. Nature, 437(7063), 1257–1263. 10.1038/nature04284

Schiff, N. D. (2010). Recovery of consciousness after brain injury: A mesocircuit hypothesis. Trends in Neurosciences, 33(1), 1–9. 10.1016/j.tins.2009.10.002

Schiff, N. D. (2023). Mesocircuit mechanisms in the diagnosis and treatment of disorders of consciousness. La Presse Médicale, 52(2), 104161. 10.1016/j.lpm.2022.104161

Schmahmann, J. D. (2019). The cerebellum and cognition. Neuroscience Letters, 688, 62–75. 10.1016/j.neulet.2018.07.005

Slot, K. B., Berge, E., Dorman, P., Lewis, S., Dennis, M., & Sandercock, P. (2008). Impact of functional status at six months on long-term survival in patients with ischaemic stroke: Prospective cohort studies. BMJ, 336(7640), 376–379. 10.1136/bmj.39456.688333.BE

Snider, S. B., Bodien, Y. G., Edlow, B. L., et al. (2020). Cortical lesions causing loss of consciousness are functionally connected to the dorsal brainstem. Annals of Neurology, 87(3), 489–494. 10.1002/ana.25678

Soddu, A., Demertzi, A., Vanhaudenhuyse, A., Bruno, M.-A., Tshibanda, L., Di, H., … Laureys, S. (2020). Resting-state activity in patients with disorders of consciousness. Functional Neurology, 35(3), 123–130. 10.11138/FNeur/2020.35.3.123

Sperber, C., & Karnath, H.-O. (2017). Impact of correction factors in human brain lesion–behaviour inference. Human Brain Mapping, 38(3), 1692–1701. 10.1002/hbm.23490

Steriade, M., & McCarley, R. W. (2005). Brain control of wakefulness and sleep. Boston, MA: Springer US.

Stoodley, C. J., & Schmahmann, J. D. (2018). Functional topography of the human cerebellum. In M. Manto (Ed.), Handbook of Clinical Neurology (Vol. 154, pp. 59–70). Elsevier. 10.1016/B978-0-444-63956-1.00005-9

Tao, W. D., Liu, M., Fisher, M., Wang, D. R., Li, J., Furie, K. L., … & Wu, B. (2012). Posterior versus anterior circulation infarction: how different are the neurological deficits? Stroke, 43(8), 2060–2065. 10.1161/STROKEAHA.112.652420

Teasdale, G., & Jennett, B. (1974). Assessment of coma and impaired consciousness: A practical scale. The Lancet, 304(7872), 81–84. 10.1016/S0140-6736(74)91639-0

Vemmos, K., Bots, M. L., Tsibouris, P. K., Zis, V. P., Takis, C. E., Grobbee, D. E., & Stamatelopoulos, S. (2000). Prognosis of stroke in the south of Greece: 1 year mortality, functional outcome and its determinants: the Arcadia Stroke Registry. *Journal of Neurology*, Neurosurgery & Psychiatry, 69(5), 595–600. 10.1136/jnnp.69.5.595

Wang, Z., Zhang, S., Liu, C., et al. (2019). A study of neurite orientation dispersion and density imaging in ischemic stroke. Magnetic Resonance Imaging, 57, 28–33. 10.1016/j.mri.2018.10.018

Warren, A. E. L., Raguž, M., Friedrich, H., et al. (2025). A human brain network linked to restoration of consciousness after deep brain stimulation. Nature Communications, 16, 6721. 10.1038/s41467-025-63046-5

Wu, O., Cloonan, L., Mocking, S. J., Bouts, M. J., Copen, W. A., Cougo-Pinto, P. T., … & Rost, N. S. (2015). Role of acute lesion topography in initial ischemic stroke severity and long-term functional outcomes. Stroke, 46(9), 2438–2444. 10.1161/STROKEAHA.115.009643

Young, G. B. (2009). Coma. Annals of the New York Academy of Sciences, 1157(1), 32–47. 10.1111/j.1749-6632.2009.04471.x

Zeng, W., Chen, Y., Zhu, Z., Gao, S., Xia, J., Chen, X., … Zhang, Z. (2020). Severity of white matter hyperintensities: Lesion patterns, cognition, and microstructural changes. Journal of Cerebral Blood Flow & Metabolism, 40(12), 2454–2463. 10.1177/0271678X1989360

Zhang, Y., Kimberg, D. Y., Coslett, H. B., Schwartz, M. F., & Wang, Z. (2014). Multivariate lesion-symptom mapping using support vector regression. Human Brain Mapping, 35(12), 5861–5876. 10.1002/hbm.22590

